# PhyloNaP: a user-friendly database of Phylogeny for Natural Product–producing enzymes

**DOI:** 10.1101/2025.09.23.677986

**Authors:** Aleksandra Korenskaia, Judit Szenei, Lisa Vader, Kai Blin, Tilmann Weber, Nadine Ziemert

## Abstract

Phylogenetic analysis is widely used to predict enzyme function, yet building annotated and reusable trees is labor-intensive and requires extensive knowledge about the specific enzymes. Existing resources rarely cover biosynthetic enzymes and lack the context needed for meaningful analysis.

We present PhyloNaP, the first large-scale resource dedicated to phylogenies of biosynthetic enzymes. PhyloNaP provides ∼18,500 annotated and interactive trees enriched with chemical, functional, and taxonomic information. Users can classify their own sequences via phylogenetic placement, enabling functional inference in an evolutionary context. A contribution portal allows the community to submit curated trees. By combining scale, breadth of annotation, and interactive functionality, PhyloNaP fills a major gap in bioinformatics resources for enzyme discovery and annotation, with immediate applications to secondary metabolism and beyond.

**Availability:** https://phylonap.cs.uni-tuebingen.de

## 1 Introduction

Phylogenetic analysis is a powerful approach for predicting enzyme function, offering insights that go beyond what can be inferred from sequence similarity alone. By placing proteins into an evolutionary context, phylogenies help delineate functional subgroups, identify catalytic mechanisms, and uncover novel biochemistry. This makes phylogeny a widely used strategy in enzyme discovery, biocatalyst engineering, and metabolic pathway reconstruction.

This challenge is particularly acute for enzymes and proteins involved in the biosynthesis of natural products, which are typically encoded in biosynthetic gene clusters (BGCs). Core biosynthetic enzymes such as polyketide synthases (PKSs) and nonribosomal peptide synthetases (NRPSs) can not only be readily detected through conserved domains, but in many cases their modular organization also enables reliable predictions of the core scaffold they assemble (Terlouw et al. 2025). By contrast, tailoring enzymes—including halogenases, glycosyltransferases, oxidoreductases, and methyltransferases—pose a different problem. Their broad enzyme class can usually be assigned from conserved sequence motifs, but it is far more difficult to predict their precise biochemical role: which substrate they act on, the position of a modification, and the type of chemical transformation they perform. These details are crucial, as tailoring reactions define the structural diversity and biological activity of the final compounds (Walsh 2023). In such cases, phylogenetic placement against experimentally characterized homologs provides a more fine-grained basis for functional inference than domain annotation alone (Adamek et al. 2019).

Several widely used genome mining tools have adopted phylogeny-based strategies to improve functional annotation. For example, NaPDoS classifies ketosynthase (KS) and condensation (C) domains based on curated phylogenetic trees to support the identification of PKS and NRPS enzymes (Ziemert et al. 2012). antiSMASH incorporates fine-scale phylogenetic models and profile Hidden Markov Models (pHMMs) to predict substrate specificities and BGC product classes (Blin et al. 2025). However, building high-quality phylogenies remains labor-intensive: it requires assembling training sets, aligning sequences, inferring trees, and annotating them with functional or chemical data. Published trees are typically static images that cannot be reused for classifying new sequences. More general phylogenetic databases, such as PhylomeDB (Fuentes et al. 2022) or EggNOG (Hernández-Plaza et al. 2023), provide broad coverage but underrepresent biosynthetic enzymes and lack integration with chemical or genomic context, limiting their utility for this application.

To address these limitations, we developed PhyloNaP, a web-based platform and database of annotated phylogenetic trees for biosynthetic enzymes. PhyloNaP currently hosts ∼18,500 trees, integrating chemical, functional, and taxonomic metadata from multiple sources. The resource supports interactive exploration of trees with customisable metadata display and phylogenetic placement of user-submitted sequences. A contribution portal allows researchers to submit manually annotated trees, ensuring continuous growth supported by the research community.

By bridging the gap between phylogenetic analysis and functional annotation, PhyloNaP provides a community-driven resource of annotated biosynthetic enzyme phylogenies that integrates chemical and functional metadata, connects to complementary databases such as MITE (Mitja M. Zdouc et al. 2024), and facilitates applications ranging from structure prediction to enzyme engineering in synthetic biology.

## 2 Database content and construction

PhyloNaP currently hosts approximately 18,500 phylogenetic trees, each representing a distinct protein family or subfamily involved in natural product biosynthesis (see Figure 1A). Each dataset comprises a rooted phylogenetic tree in Newick format, a multiple sequence alignment, a FASTA file of untrimmed sequences, and a metadata file with functional, taxonomic, and/or chemical annotations. The majority of sequences are of bacterial origin, with the function ranging from scaffold-forming synthases to diverse tailoring enzymes and other proteins that can be found in BGCs, such as transporters, regulatory and resistance proteins (see Figure 1B). Where available, datasets are linked to experimentally characterized reactions and validated BGCs, allowing chemical structures and reaction information to be visualized directly on the tree. For the many proteins that remain uncharacterized, PhyloNaP adds value by providing contextual information— such as BGC type and taxonomy—that guides hypothesis generation and exploratory analysis. Cross-links to external resources further connect each entry to the broader bioinformatics ecosystem and will grow richer as experimental data accumulates.

**Figure 1.**
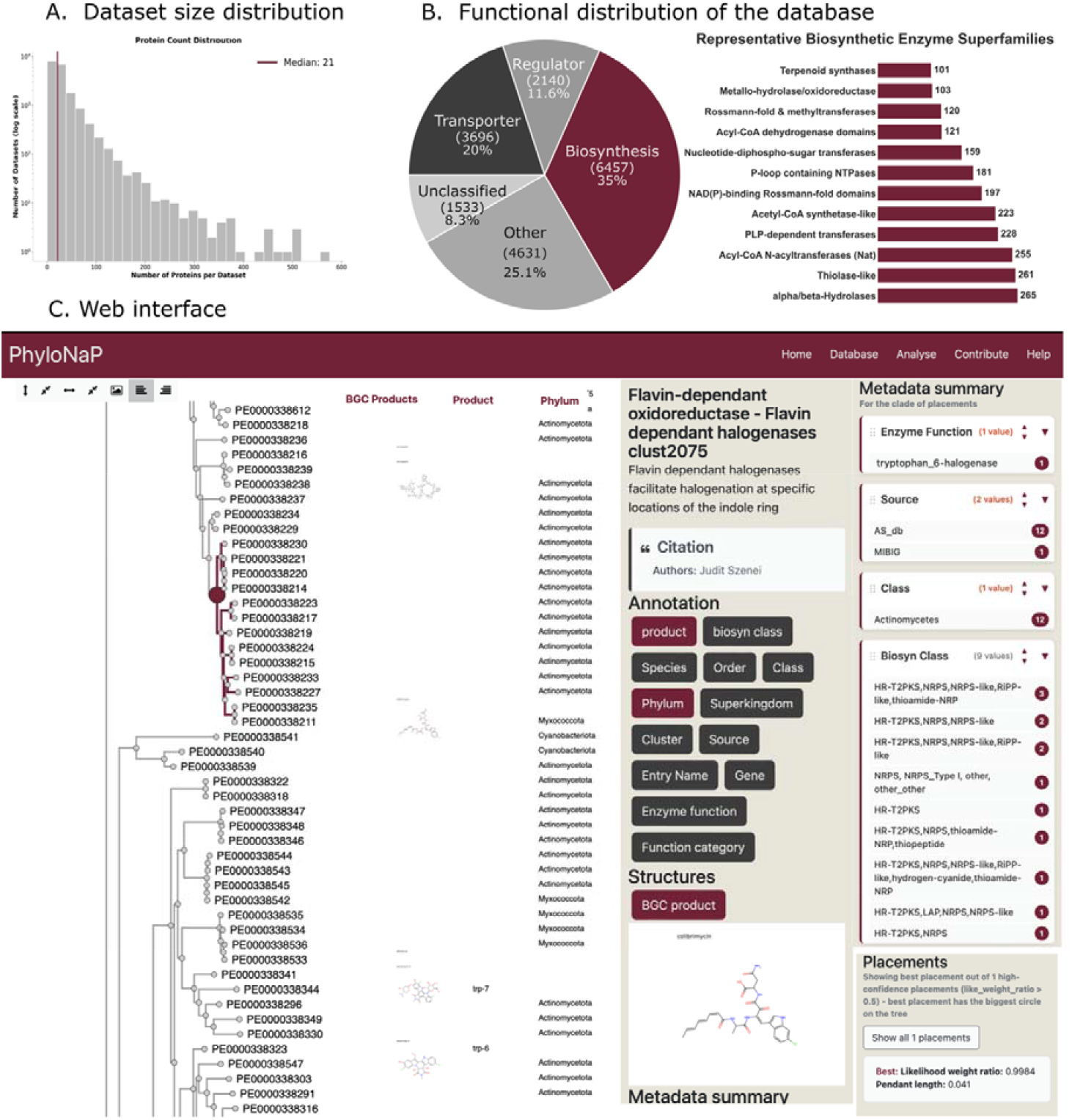
A.Dataset size distribution. B. Functional overview obtained with eggNOG-mapper (Cantalapiedra et al. 2021; Hernández-Plaza et al. 2023); see details in the Supplementary Methods. C. Tree viewer interface of PhyloNaP.

### Curated and automated datasets

PhyloNaP distinguishes between curated and automatically generated datasets. The curated datasets comprise ten enzyme families that have been manually assembled and annotated by experts, many of them derived from published studies (Reitz et al. 2019; Zdouc et al. 2021; Biermann et al. 2024; Gavriilidou et al. 2023). These serve as high-confidence reference trees for functional classification.

The majority of datasets are generated through an automated pipeline (see Supplementary Methods) that integrates protein sequences and annotations from established reference databases, clusters them into families (Steinegger and Söding 2018), and then consistently aligns (Katoh and Standley 2013), trims (Capella-Gutiérrez et al. 2009), and infers rooted trees (Price et al. 2010; Bryant and Charleston 2018) suitable for exploration and sequence placement. Each dataset is further classified into enzyme superfamilies using HMMER with Superfamily 4.0 profiles (Pandurangan et al. 2019). This workflow enables large-scale coverage while maintaining consistent quality. With ongoing community contributions of curated trees, PhyloNaP is designed to evolve into a comprehensive resource that combines expert knowledge with automated scalability. Ongoing community submissions will steadily expand the number of curated families, ensuring PhyloNaP continues to evolve as a reference resource.

## 3 Classifying user sequences

A key feature of PhyloNaP is its ability to classify user-submitted sequences by placing them onto existing phylogenetic trees. This workflow allows researchers to interpret their sequences in an evolutionary context and infer likely functions or biochemical roles.

Users begin by submitting one or more protein sequences in FASTA format. The system first uses MMseqs2 (Steinegger and Söding 2017) to identify similar proteins within the PhyloNaP database, using a default minimum threshold of 30% identity and 50% coverage. For each matching reference tree, the query sequence is added to the corresponding alignment, and phylogenetic placement is performed (Barbera et al. 2019), usually within seconds or minutes, to determine the most likely positions of the query sequence along the tree.

## 4 Web Interface and User Tools

We designed PhyloNaP with a strong emphasis on usability, creating an interface that allows researchers to both explore the database and analyze their own sequences in a single environment. The entry point is a database browser, where users can search and filter thousands of annotated phylogenies by enzyme family, dataset origin (curated or automated), and annotation type. This makes it straightforward to identify relevant reference trees for a particular research question.

To support functional inference, PhyloNaP offers a dedicated Analyze page where users can upload protein sequences in FASTA format. The system automatically identifies the most similar datasets, performs phylogenetic placement, and reports results in a concise table. Each entry provides a description of the matched dataset, quantitative placement scores such as the likelihood weight ratio (LWR)(Matsen et al. 2012), and direct links to an interactive tree viewer.

The interactive viewer is central to PhyloNaP. It provides dynamic exploration of phylogenetic trees with features such as rerooting, zooming, and adjustable branch spacing. Multiple layers of metadata, including functional predictions, taxonomic information, BGC classifications, and, when available, chemical structures or reaction schemes, can be overlaid directly on the tree (Figure 1C). These annotations are fully interactive. Structures can be expanded for inspection, and hyperlinks lead directly to external resources such as MiBiG (Mitja M Zdouc et al. 2024), which links experimentally validated BGCs to their products; SwissProt (Boutet et al. 2007; The UniProt Consortium 2025), providing curated protein sequences with functional data; MITE (Mitja M. Zdouc et al. 2024), which contributes manually curated reaction data for tailoring enzymes; and PanBGC (Paccagnella et al. 2025) enabling comparative analysis of similar BGCs from the antiSMASH database (Blin et al. 2024). Together, these links provide users with direct access to high-quality chemical, functional, and genomic context. When user-submitted sequences are placed, they appear as highlighted nodes with confidence scores visualized by circle size. A metadata summary of the most recent common ancestor of all placement nodes is displayed by default, providing an immediate overview of the functional and genomic context.

Beyond browsing and analysis of phylogenetic trees, PhyloNaP also supports community engagement. A submission portal enables researchers to contribute their own curated datasets, which are reviewed for quality before being integrated into the public resource. This feature ensures that the database continues to grow not only in scale but also in depth, combining expert knowledge with automated coverage.

As an illustration, we tested the placement of a flavin-dependent halogenase (A0A1L1QK36 in UniProt) (Figure 1C). The analysis page reports placements onto two reference datasets, with the prioritized tree showing a high LWR (0.998) and a short branch length (0.04). The sequence is positioned in a clade dominated by proteins from NRPS BGCs, suggesting amino acid or peptide substrates. One closely related enzyme is annotated as a tryptophan-6 halogenase, and the natural product structures linked to this clade consistently carry a halogen substituent at this position.

Taken together, the reliable positioning, very short branch length, and phylogenetic and chemical evidence strongly suggest that the query protein performs a tryptophan-6 halogenation This example demonstrates how PhyloNaP’s interface integrates evolutionary, functional, and chemical information to support detailed functional predictions; a more detailed description of the workflow is provided in the Supplementary Methods.

## 5 Conclusion and discussion

PhyloNaP establishes a centralized platform for annotated phylogenetic trees of biosynthetic enzymes. By integrating functional, chemical, and taxonomic metadata into interactive phylogenies, it enables researchers to investigate enzyme families in an evolutionary framework and to classify novel sequences with confidence.

A key strength of PhyloNaP lies in its integration with external resources such as MiBiG, MITE, SwissProt, and PanBGC. These links enrich the evolutionary trees with experimentally validated functions, reaction data, and genomic context, allowing users to connect sequence placement directly to biochemical and structural information. Such integration facilitates practical applications ranging from structure prediction of natural products to the identification of candidate enzymes for pathway engineering in synthetic biology.

The resource is not without limitations. Automated tree rooting is based on minimal ancestor deviation and may be suboptimal for some families, although users can interactively reroot trees for visualization. Functional annotations depend on the current state of reference databases, and many biosynthetic enzymes remain poorly characterized, meaning that predictions often reflect context rather than exact reaction specificity. In addition, sequence placement is currently limited to 50 sequences to ensure computational efficiency, and manual inspection remains essential for reliable interpretation.

Looking ahead, PhyloNaP is designed as a living, community-driven resource. Regular updates to MiBiG and MITE will steadily expand the pool of experimentally characterized proteins, while direct user contributions will add expert-curated phylogenies. Future development will focus on tighter integration with genome-mining and machine-learning tools, deeper curation of submitted datasets, and expanded coverage of published trees. By reducing redundancy, improving accessibility, and fostering community input, PhyloNaP has the potential to accelerate functional annotation, enable discovery of new enzymatic activities, and support the rational exploration of biosynthetic diversity.

## Data availability

The PhyloNaP database is available for browsing and downloading individual datasets at https://phylonap.cs.uni-tuebingen.de/database, and the full database can be downloaded from https://phylonap.cs.uni-tuebingen.de/download. The results of the functional distribution analysis reported in this article are provided as Supplementary Table. The source code of the PhyloNaP web application is available at https://github.com/ZiemertLab/PhyloNaP_WebApp.

## Acknowledgments

We would like to acknowledge Noel Kubach, Dr. Mitja Zdouc, Mathiaz Witte Paz, and Konstantin Vysotskii for discussions and advice on the tool development; and Dr. Leo Padva and Dr. Athina Gavriilidou for sharing datasets. Artificial intelligence (AI) tools (GPT-4o and GPT-5, OpenAI; Claude Sonnet 4, Anthropic) were used to assist with text editing and code writing and debugging.

