## Supplementary Methods for "PhyloNaP: a user-friendly database of Phylogeny for Natural Product–producing enzymes"

### Dataset generation workflow

The dataset generation pipeline consists of a series of automated steps designed to collect, filter, and organize protein sequences into phylogenetically structured datasets.

Protein sequences were collected from several established resources. The MiBiG 4.0 database (Mitja M Zdouc et al. 2024) provided proteins from experimentally characterized biosynthetic gene clusters (BGCs), while MITE (Mitja M. Zdouc et al. 2024) contributed enzymes with experimentally validated reactions. AntiSMASH-DB 4.0 (Blin et al. 2024) supplied predicted BGC-derived proteins from large-scale genome mining, and UniProt SwissProt (Boutet et al. 2007; The UniProt Consortium 2025) added manually curated sequences with high-quality functional data.

Each sequence was enriched with metadata. Taxonomic information was retrieved via the NCBI Datasets Taxonomy Data Package (O’Leary et al. 2024). Functional annotations, BGC product classes, and cross-links to external databases were added where available. Structural data for substrates or products were obtained either directly from MITE and MiBiG or indirectly via UniProt cross-references to ChEBI (Hastings et al. 2016) and Rhea (Bansal et al. 2022). RDKit (Landrum et al. 2020) was used to generate chemical structure depictions. Sequences from antiSMASH-DB were additionally mapped to PanBGC (Paccagnella et al. 2025) to enable cross-referencing.

Sequences were clustered into datasets using MMseqs2 (*easy-linclust*) (Steinegger and Söding 2017) with an e-value threshold of 1e-3. Several filtering steps were applied to ensure biological relevance and computational feasibility. Clusters with fewer than 20 sequences, clusters containing only eukaryotic sequences or only SwissProt entries (probably primary metabolism), clusters with extreme mean sequence lengths (<150 or >1000 amino acids), and clusters dominated by NRPSs or PKSs were excluded, as these modular enzymes require domain-level analyses.

For each retained cluster, multiple sequence alignment was performed with MAFFT (auto mode) (Katoh and Standley 2013), followed by automatic trimming with TrimAl (Capella-Gutiérrez et al. 2009). Identical sequences were removed after trimming, while unique annotation information was retained in dedicated metadata fields in columns <original column name>_others. Clusters reduced to fewer than 10 sequences after this step were discarded. Phylogenetic trees were inferred using FastTree (Price et al. 2010) and consistently rooted with MADroot (Bryant and Charleston 2018). While curated datasets rely on evolutionary models specifically selected for each case, large-scale automated inference requires a faster and more uniform approach. FastTree, which is well suited for generating thousands of trees efficiently, applies the JTT (Jones–Taylor–Thornton) model of amino acid substitution by default. Accordingly, all automatically generated datasets were inferred under the JTT model, ensuring consistency and comparability across the collection.

Each dataset was assigned to an enzyme superfamily based on HMMER searches against the Superfamilies v1.75 (Pandurangan et al. 2019) profile set. All detected domains were aggregated to generate a dataset-level superfamily annotation. Datasets with identical superfamily combinations were grouped together. All datasets and associated metadata were stored both as a structured JSON index and in a NoSQL database, enabling efficient access and filtering. For each dataset, summary statistics were compiled, including the total number of sequences, the number of experimentally characterized enzymes (from SwissProt with Rhea annotations or from MITE), the number of proteins derived from validated BGCs (MiBiG), and the number of proteins from predicted BGCs (antiSMASH-DB).

This procedure results in a collection of high-quality, phylogenetically structured datasets optimized for functional exploration and user-guided sequence placement.

### Database functional overview

The functional diversity of datasets (Figure 1B in the article) was assessed in two complementary ways.

**Functional categories from eggNOG.**

To provide an overview of functional diversity, one representative sequence was selected from each dataset. The representative was defined as the central sequence, i.e. the sequence with the minimal sum of evolutionary distances to all other sequences in the phylogenetic tree. Representatives were annotated using eggNOG-mapper (Cantalapiedra et al. 2021) using EggNOG 6.0 database (Hernández-Plaza et al. 2023) assign general functional roles.

Datasets were classified into five major categories based on COG functional assignments: biosynthetic, transporter, regulator, other, and unclassified. Proteins associated with amino acid, nucleotide, lipid, coenzyme, energy, or secondary metabolite metabolism were classified as biosynthetic. Transporter proteins included those linked to inorganic ion or carbohydrate transport. Sequences annotated under transcription or signal transduction were assigned to the regulator group. Proteins with functions related to primary metabolism, resistance, secretion, or general cellular processes were classified as other, while sequences without a confident assignment remained unclassified. To improve transporter identification and prevent misclassification, the protein description field was screened for transporter-related terms, including “ABC transporter”, “major facilitator”, “MFS”, “permease”, “transporter”, “channel protein”, “importer”, and “exporter”. Matches were reclassified as transporters when appropriate.

When multiple COG categories were assigned to a single sequence, sequence were classified to one of the functional groups followed a hierarchical rule: transporter > regulator > biosynthesis > other > unclassified. This procedure ensured consistent categorization across datasets.

**Superfamily-based overview.**

|  | **Hmm combination** | **Dataset count** | **General function** |
| --- | --- | --- | --- |
| **1** | MFS general substrate transporter | 461 | Transporter |
| **2** | P-loop containing nucleoside triphosphate hydrolases\|PEP carboxykinase-like | 399 | Other |
| **3** | Winged helix DNA-binding domain\|Periplasmic binding protein-like II | 284 | Regulator |
| **4** | alpha/beta-Hydrolases | 265 | Biosynthetic |
| **5** | Thiolase-like | 261 | Biosynthetic |
| **6** | Acyl-CoA N-acyltransferases (Nat) | 255 | Biosynthetic |
| **7** | MetI-like | 236 | Transporter |
| **8** | PLP-dependent transferases | 228 | Biosynthetic |
| **9** | Acetyl-CoA synthetase-like | 223 | Biosynthetic |
| **10** | Periplasmic binding protein-like II | 218 | Transporter |
| **11** | NAD(P)-binding Rossmann-fold domains | 197 | Biosynthetic |
| **12** | P-loop containing nucleoside triphosphate hydrolases | 181 | Biosynthetic |
| **13** | Nucleotide-diphospho-sugar transferases | 159 | Biosynthetic |
| **14** | Porins | 143 | Transporter |
| **15** | ClpP/crotonase | 135 | Other |
| **16** | FAD/NAD(P)-binding domain\|MurCD N-terminal domain\|NAD(P)-binding Rossmann-fold domains\|Nucleotide-binding domain | 132 | Other |
| **17** | Acyl-CoA dehydrogenase C-terminal domain-like\|Acyl-CoA dehydrogenase NM domain-like | 121 | Biosynthetic |
| **18** | NAD(P)-binding Rossmann-fold domains\|S-adenosyl-L-methionine-dependent methyltransferases | 120 | Biosynthetic |
| **19** | ABC transporter involved in vitamin B12 uptake\|BtuC | 118 | Transporter |
| **20** | Metallo-hydrolase/oxidoreductase | 103 | Biosynthetic |
| **21** | Terpenoid synthases | 101 | Biosynthetic |

Supplementary Table 1. Largest enzyme superfamilies represented in the PhyloNaP database. Dataset count shows the number of datasets within the superfamily.

In parallel, the superfamily annotations previously obtained during the dataset-generation stage (via HMMER searches against the Superfamily v1.75 profiles) were retrieved and used to highlight dominant enzyme families. Datasets sharing identical superfamily assignments were grouped, and superfamilies represented by more than 100 datasets were manually inspected. Their functional roles were then assigned into the same five major categories used above. The largest biosynthetic superfamilies are shown in Figure 1B (right panel). A complete list of the top 21 superfamilies, together with their dataset counts, is provided in Supplementary Table 1.

### A workflow example

As an illustration, we tested the placement of a flavin-dependent halogenase (A0A1L1QK36 in UniProt). The analysis page shows placements onto two reference datasets (Supplementary figure 1). By default, only the best-scoring dataset is displayed, but the user can toggle a button to view alternatives. Within a tree, the query sequence may be placed onto a leaf or an internal node, and sometimes multiple placement options are reported. Typically, the more divergent a query sequence is from the reference tree sequences, the more alternative placements are reported, reflecting increased uncertainty (Matsen et al. 2012; Barbera et al. 2019).

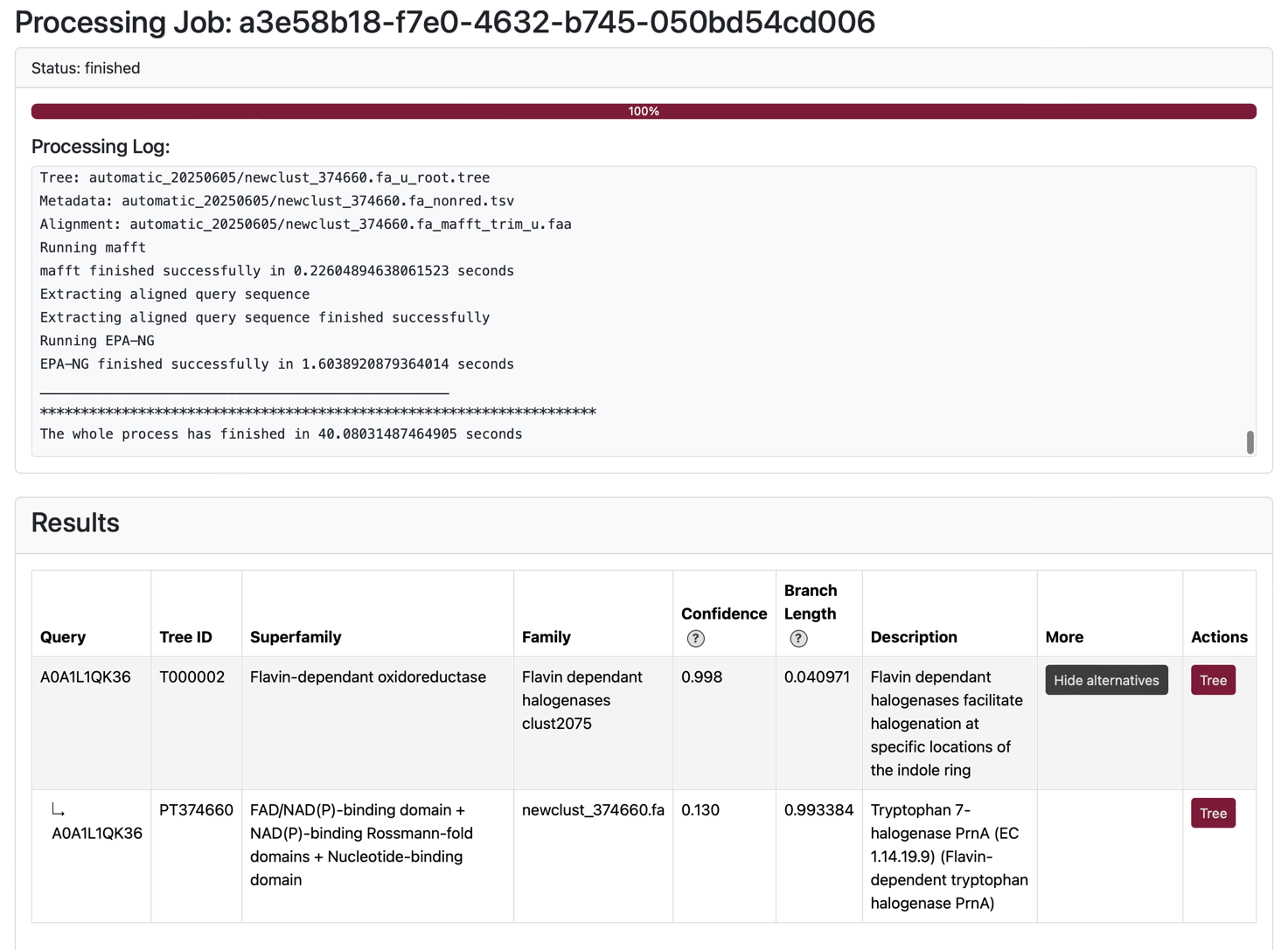

Supplementary Figure 1. Query sequence placements on two reference datasets.

The **confidence column** reports the likelihood weight ratio (LWR) of the best placement for each reference dataset. LWR measures how strongly the data support a given placement relative to all other possible placements of the same query sequence within that dataset. All LWRs for the query within one tree sum to 1. In our example, the placement in the prioritized reference dataset has an LWR of 0.998, while the best placement in the second reference dataset has an LWR of 0.130, indicating that the first dataset provides much stronger support.

The **branch length** indicates the evolutionary distance between the query and the node of placement based on the alignment and substitution model. A short branch length suggests closer similarity, which makes it more reasonable to infer function from the surrounding clade. Conversely, a long branch length suggests that the alignment places the query far from its nearest reference, and any inference becomes more tentative. In this case, the placement on the prioretised reference dataset has a short branch length (≈0.04 substitutions per site), supporting a reliable placement, while in the second reference dataset the branch length is nearly 1, signaling much weaker support.

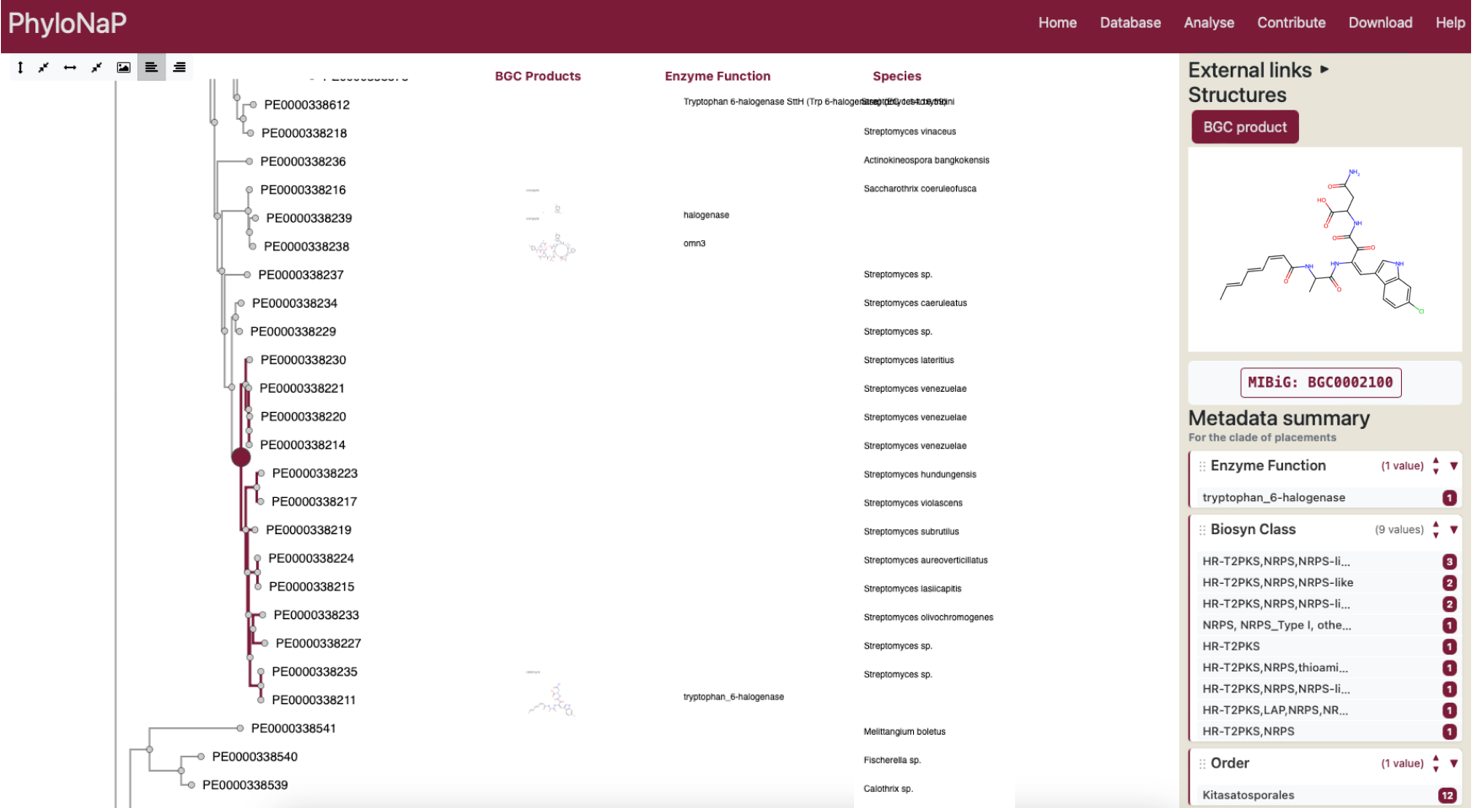

Supplementary Figure 2. Placement node (red circle) and corresponding clade (highlighted in red) with Annotations and Metadata summary.

Let us now inspect the dataset with the reliable placement. The placement node is marked in red circle, and the clade defined by this node is highlighted in the same color (Supplementary Figure 2). All available annotations for this clade are summarized in the “Metadata summary” field, which shows how consistent each type of annotation is. From this, we see that nearly all proteins in the clade, except one, are associated with nonribosomal peptide synthetase (NRPS) biosynthetic gene clusters, indicating that their substrates are amino acids or peptides. All proteins in the clade come from species of the genus Streptomyces. One enzyme carries a functional annotation as a tryptophan-6 halogenase.

To investigate further, we can click the “Enzyme function” button in the metadata panel to display this annotation directly. Additionally, the “Structures” field provides the option to visualize BGC products: by selecting it, images of natural products from the MIBiG database, in whose biosynthesis the proteins are involved, appear. In this case, one protein in the clade has both a functional annotation and a linked product structure, allowing us to cross-validate the assignment and confirm that it is indeed a tryptophan-6 halogenase.

A.

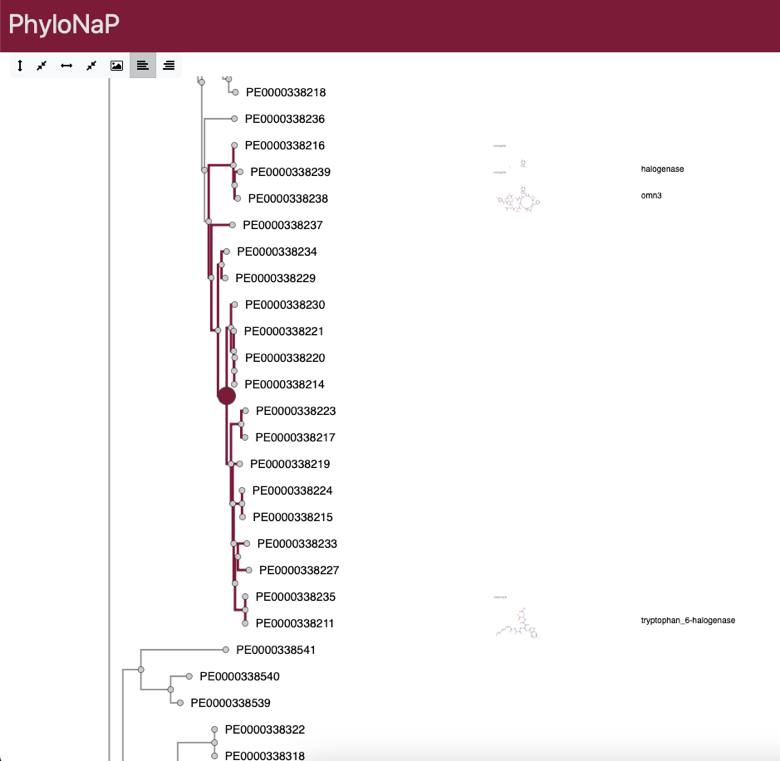

B.

.
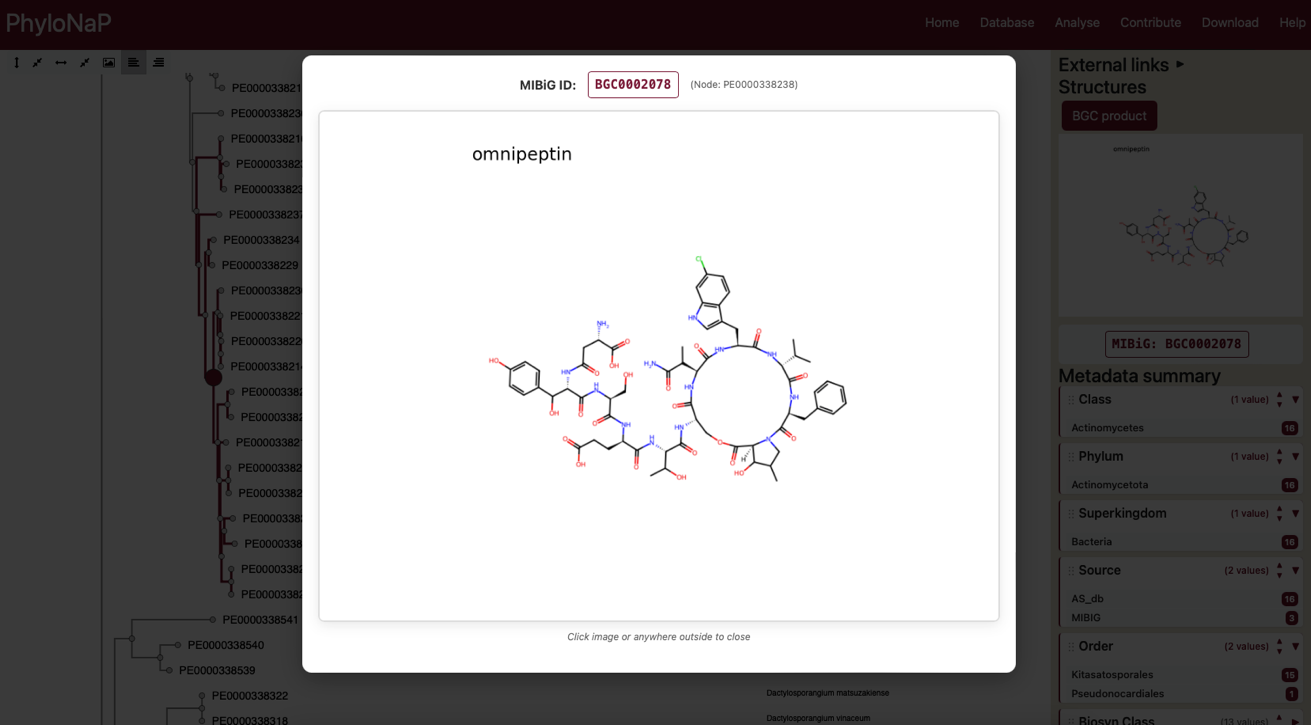

**Supplementary Figure 3.** A. Smallest ancestral clade including additional annotated proteins (highlighted in red). B. Enlarged chemical structure of a natural product associated with an additional annotated protein, showing a halogen substituent at position 6 of tryptophan.

Since no other detailed annotations are available within the clade itself, we can examine the nearest annotated ancestral node (Supplementary Figure 3A), the clade is indicated in red. Inspection of the proteins branching from this node reveals another natural product that clearly carries a halogen substituent at position 6 of tryptophan (Supplementary Figure 3B). Even though the text annotation of the associated enzyme function does not specify the substitution site, the chemical structure image confirms it.

In summary, the reliable placement, the short branch length, and the agreement between phylogenetic and chemical evidence all point to the same conclusion: the query protein most likely carries out the same reaction. The clade, defined by their common ancestor, shows consistent functional annotations, supporting that our enzyme catalyzes halogenation of tryptophan at position 6. This combination of visual and contextual information enables researchers to infer substrate class and biochemical role, guiding downstream functional studies.
